# Impact of Mesotherapy with Sodium Deoxycholate on Liver: Metabolic- and Sex-Specific Insights in Swiss mice

**DOI:** 10.1101/2024.01.18.576283

**Authors:** Leidyanne Ferreira Gonçalves, Beatriz Rodrigues Rosa, Isabela Terra Tavares Ramos, Julia Bueno Feder, Julia Rajczuk Martins Messina, Raissa Moreira Barreira, Vanessa Morales Torres, Vitor Lima Simões, Elan Cardozo Paes-de-Almeida, Caroline Fernandes-Santos

**Author notes:** Corresponding Author: Rua Dr. Silvio Henrique Braune, 22, Centro, Nova Friburgo, RJ, Brazil, 28.625-650. Funding source: Fundação Carlos Chagas Filho de Amparo à Pesquisa do Estado do Rio de Janeiro FAPERJ E-26/010.001892/2019. The authors have no conflict of interest to declare.

## Abstract

**Background:** Sodium deoxycholate (DC) is often used in mesotherapy for the aesthetic improvement of body contouring. Although it is a minimally invasive procedure, DC use is off-label since, to date, it is approved solely for submental fat reduction, lacking evidence to support its safety to other body regions.

**Objective:** To investigate the systemic and hepatic effects of the prolonged use of DC in mesotherapy for fat reduction in Swiss mice under fructose consumption.

**Methods:** Female and male Swiss mice received water or 20% fructose (F) ad libitum for 12 weeks. DC 50 μg sc. was administered into the right inguinal white adipose tissue (riWAT) twice weekly for 4 weeks starting week 8. We assessed body weight (BW), glucose, lipolysis, hepatic enzymes, adipose tissue remodeling, liver histopathology, and protein expression.

**Results:** Chronic DC did not affect BW, glucose, lipolysis, and hepatic enzymes, except for ALT in males. Although the riWAT weight remained stable, we found foam cells, tissue hemorrhage, and fibrosis. DC induced neither hepatomegaly nor hepatocyte hypertrophy in either sex except for fructose in females, which led to heavier livers and increased hepatocyte nuclei volume. Mild fat deposition was present in fructose-fed female mice, with no influence of DC injections. Finally, FXR and FGF21 protein expression were similar among the groups.

**Conclusion:** DC had no impact on BW or adipose tissue mass, although there were features of chronic riWAT inflammation. It failed to impair glucose and hepatic metabolism, morphology, and protein expression in both sexes.

## 1. Introduction

Mesotherapy, also called local intradermal therapy, was initially introduced in 1952 by a French doctor named Pistor [1]. The technique promotes direct transepidermal transport of active substances into the dermis and deeper skin layers [2]. It has gained increasing popularity in aesthetics for localized fat removal since it is less invasive and expensive than liposuction [3]. Sodium deoxycholate (DC), or deoxycholic acid, is a secondary bile acid produced by the gut microbiota through cholic acid metabolism that emulsifies dietary fats, allowing its absorption [4, 5]. In the United States, the use of DC has been approved since 2015, and until 2022, it was indicated solely for submental fat reduction in adults [6]. Likewise, Canada, Australia, Europe, and South Korea have approved DC use only for submental fat reduction [7]. However, DC is widely used off-label in other body regions, such as the bra line and for body contouring [8]. Consequently, studies need to be carried out to prove its safety, but only some ongoing clinical trials are investigating the metabolic impact of abdominal fat reduction by DC. Published data so far evaluated serum lipid and adipokine profile [9], DC safety and tolerability [10], and abdominal fat histopathology [11] without taking into account DC systemic effects in other organs.

Firstly, the potency of DC as a detergent on adipocytes made it the primary substance for fat reduction in mesotherapy. It permeates the adipocyte membrane, destabilizing and disintegrating the lipid bilayer, subsequently forming micelles composed of detergent and cell membrane lipids [12]. It results in fat necrosis due to the leakage of the cytoplasmic content and the massive release of triglycerides (TG) into the interstitium [13], promoting macrophage recruitment to remove lipids and cellular debris [14] and local inflammation [11]. Nevertheless, it remains uncertain whether the lipids released reach the liver, a central organ in lipid homeostasis [15]. Under normal circumstances, the liver stores small amounts of TG in cytoplasmic lipid vesicles. However, in scenarios of excessive circulating TG, such as overnutrition, obesity, and adipose tissue inflammation, the hepatic fatty acid metabolism is impaired, leading to hepatocyte TG accumulation [16]. Overall, we hypothesize that DC promotes an overflow of TG from adipose cells to the liver, leading to hepatic TG storage and tissue dysfunction.

A second issue is that DC has tissue receptors in organs enrolled in glucose and lipid metabolism [17–19]. In humans, it has been proved that DC reaches the bloodstream after subcutaneous injection. Walker et al. (2015) showed that DC 100 mg sc increases DC serum levels within the endogenous normal range, and after 12 hours, it returned to basal levels; thus, DC might have a systemic action through its receptors [9]. Moreover, DC binds to several tissues that express its receptors and may be implicated in a series of metabolic diseases [17, 20]. With this in mind, we hypothesize that the chronic use of DC in mesotherapy modulates the hepatic metabolism and gene expression through the farnesoid X receptor (FXR) [21]. Finally, the fibroblast growth factor 21 (FGF21), which connects the liver with the adipose tissue and prevents hepatic lipid deposition, might also be impacted by DC mesotherapy [22].

Overall, we investigated the systemic and hepatic effects of the prolonged use of DC in mesotherapy for fat reduction and its potential impact on hepatic fat metabolism. Male and female Swiss mice were studied under fructose consumption to investigate the role of sex and diet on the outcomes under investigation.

## 2. Materials and methods

### 2.1. Ethics

The Ethics Committee for Animal Use (CEUA) of the Universidade Federal Fluminense (UFF) approved the protocol (CEUA/UFF 1011/2017; 4232071220/2021). The investigation respected the Guide for the Care and Use of Laboratory Animals [23] and the Brazilian Guide for the Production, Maintenance, or Use of Animals in Teaching or Research Scientific Research Activities [24].

### 2.2. Animals

Female and male Swiss mice were provided by the Central Animal Care Facility (UFF, Niteroi, RJ, Brazil). They were maintained in the Animal Care Facility of the Instituto de Saude de Nova Friburgo (UFF, Nova Friburgo, RJ, Brazil) and housed under controlled temperature (21±1°C), humidity (60±10%), light/dark cycle (12 h/12 h lights on at 7 a.m.). They had free access to food and water. Water and food intake were recorded daily, and body weight (BW) was recorded weekly.

### 2.3. Study design

Twelve-week-old female (n=29) and male (n=30) Swiss mice received 20% fructose (F, fructose(D) P.A, Labsynth, Diadema, SP, Brazil) or filtered water for eight weeks. Through the following four weeks, they also received subcutaneous injections of saline (100 µl) or 50 µg DC (Sigma, D6750) in 100 µL saline twice weekly on the right inguinal white adipose tissue (riWAT) depot. In awake mice fasted for six hours, a small blood drop was obtained from the tail tip and used for glucose assessment (ACCU-Chek Performa, Roche, Jaguaré, SP, Brazil). Then, mice were anesthetized (100 mg/kg ketamine and 10 mg/kg xylazine ip), blood was collected by cardiac puncture, centrifuged, and the serum was stored at −20 °C. The liver and riWAT were harvested, weighed (Shimadzu AUX220, Kyoto, Japan), and fixed in 4% buffered formalin or stored at −80 °C. The right and left genital (gWAT) and inguinal (iWAT) fat depots were weighed to assess adiposity index, calculated as *gWAT + iWAT/BW*, expressed as %.

### 2.4. Blood and tissue biochemistry

Serum TG (K117, Bioclin, MG, Brazil), glycerol (K015/K117, Bioclin, MG, Brazil), albumin (Interkit, Belo Horizonte, MG, Brazil), and direct bilirubin (Interkit, Belo Horizonte, MG, Brazil) were analyzed by colorimetric assays, and alanine aminotransferase (ALT), aspartate aminotransferase (AST) and gamma-glutamyl transferase (GGT) (Interkit, Belo Horizonte, MG, Brazil) by kinetic assays. The liver (∼30 mg) was homogenized in 1 mL isopropyl alcohol (A1078.01.BJ, Labsynth, Diadema, SP, Brazil), centrifuged at 1.132 x g (NT805, Nova Técnica, Piracicaba, SP, Brazil), and the supernatant reserved for hepatic TG and glycerol assays. Assays were adapted for use in a 96-well microplate, and the wavelength was read in a microplate spectrophotometer (Epoch^TM^, BioTek, Vermont, USA).

### 2.5. Qualitative and quantitative histopathology

Liver and riWAT samples followed the routine histological processing for formalin-fixed paraffin-embedded tissues, sectioned at 5 µm thick, and stained with hematoxylin and eosin (H&E) and Masson’s trichrome. The riWAT was investigated qualitatively for fibrosis and inflammation. Liver hepatocyte nuclei density and hypertrophy were investigated, respectively, by the number density of hepatocyte nuclei (*Q*_*A*[*nuclei*]_) and the volume-weighted mean nuclear volume 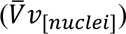 [25]. Hepatic steatosis was analyzed by pointing counting (volume density, *Vv*_[*lipid droplets*]_) in frozen liver samples embedded in OCT, sectioned at 10 µm thick, and stained with Oil red O [26, 27]. Digital images were acquired on a Leica DM750 microscope and an ICC50H video camera (Heerbrugg, Switzerland).

### 2.6. Western blot

Total proteins were extracted from ∼150-200 mg liver (RIPA lysis buffer). Proteins were separated by SDS-PAGE (10 % polyacrylamide gel) and transferred to a PVDF membrane (RNP303F, GE Healthcare, Little Chalfont, Buckinghamshire, UK) in a semi-dry blotter (TE70X, Amershan Hoefer, Holliston, MA, USA). Membranes were incubated in 5% non-fat dry milk for 1 h and then overnight with primary antibody anti-FGF21 (1:1,000, rabbit, SAB2108136, Sigma), anti-FXR (1:1,000, mouse, 72105S, Cell Signaling), and anti-β-actin (internal control, 1:5,000, mouse, sc-2005, Santa Cruz). The following day, membranes were washed with T-TBS, incubated in anti-rabbit (1:2,500, PI-1000-1, Vector Labs) or anti-mouse (1:2,500, PI-2000-1, Vector Labs) secondary antibodies, and the reaction was visualized by ECL (ThermoFisher, Pierce™ ECL Western Blotting Substrate, Carlsbad, CA, USA) using a chemiluminescence analyzer (c-DiGit Blot Scanner, LI-COR). The Image Studio Lite Software v. 5.0 (LI-COR) quantified the relative optical density.

### 2.7. Statistical analysis

Data are presented as mean ± standard deviation (S.D.). The Shapiro-Wilk test analyzed data distribution and variance homoscedasticity. The two-way ANOVA analyzed the effect of the independent factors DC injection and fructose consumption and its interaction on dependent variables, and the Holm-Sidak post hoc test was used for multiple comparisons (p<0.05, GraphPad Prism v. 8 for Windows, Boston, MA, USA).

## 3. Results

### 3.1. DC-induced inflammation and fibrosis in riWAT without fat or body weight loss

In females, fructose increased BW progressively, but not in males (Fig. 1A). On the other hand, chronic use of DC had no impact on the BW of female and male mice in the DC and F/DC groups, compared to C and F groups, respectively (Fig. 1A). Similarly, BW variation (ΔBW) during the treatment period (8-12 weeks) did not show any change in either males or females (Fig. 1B). Likewise, the adiposity index was not affected by DC (Fig. 1C), as well as the riWAT mass, which was the site of DC administration (Fig. 1D). The histopathological analysis showed adipocytes with a regular appearance in the riWAT of C and F female and male mice (Fig. 1E). In contrast, mice receiving DC (DC and F/DC groups) presented features of tissue hemorrhage, fibrosis indicating chronic injury, and foam cells which represent macrophages phagocyting adipocyte debris as crown-like structures (Fig. 1E).

**Figure 1.**
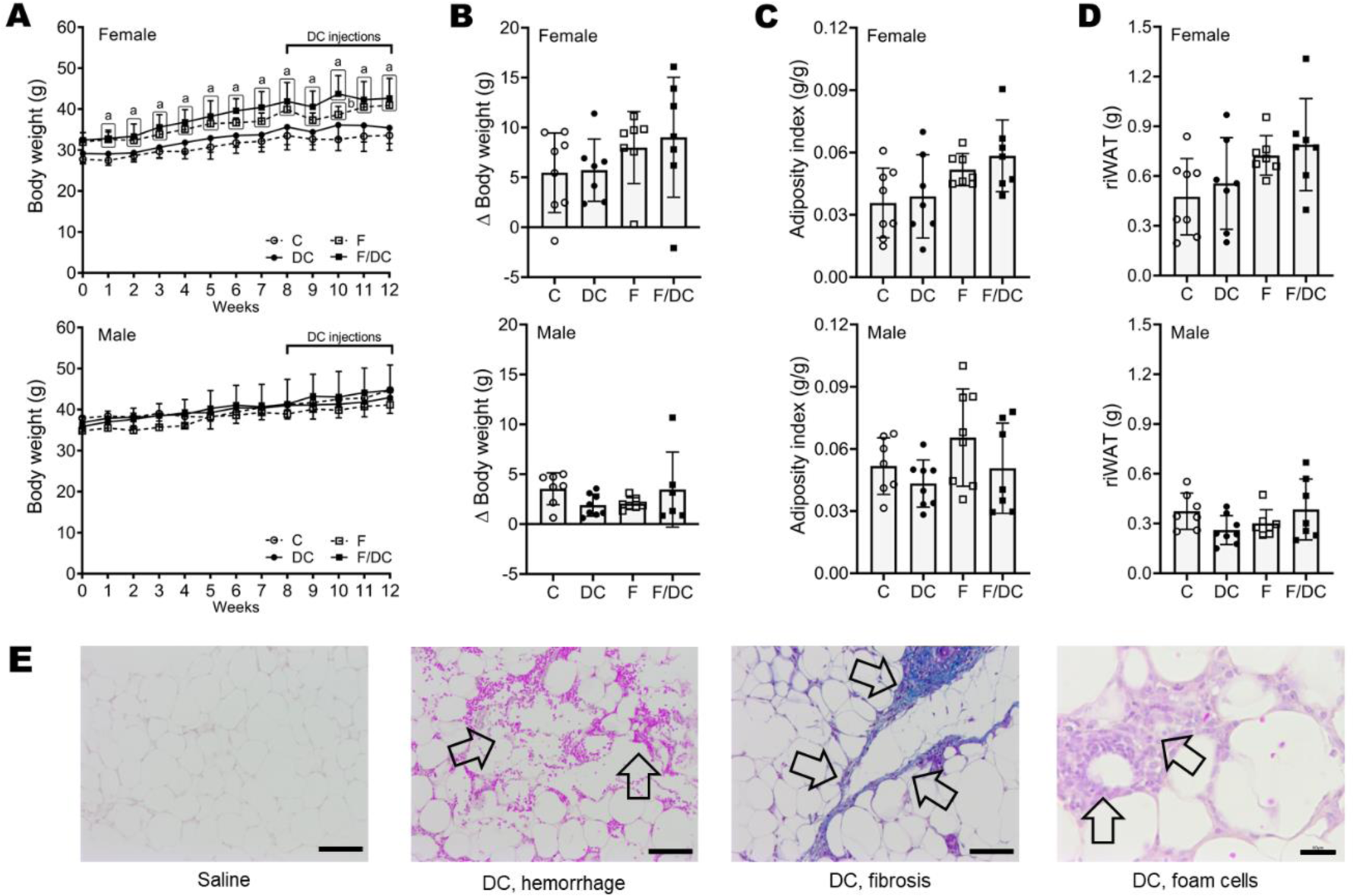
– Effect of subcutaneous injections of sodium deoxycholate (DC) on body weight and adiposity in female and male Swiss mice. **A**, Body weight. **B**, Body weight gain from weeks 8 to 12. **C**, Adiposity index. **D**, Right inguinal white adipose tissue (riWAT) weight. **E**, Representative photomicrographs of riWAT. The first image on the left shows adipocytes in saline-injected female C mice. The following images represent common DC-induced tissue injury (arrows): hemorrhage (HE stain), fibrosis (Masson’s trichrome stain), and foam cells as crown-like structures (HE stain) in female DC. Data are shown as mean ± S. D., p<0.05, [a] vs. C, [b] vs. F. Abbreviations: C, control; DC, sodium deoxycholate; F, fructose; F/DC, fructose + sodium deoxycholate. Bar = 60 µm.

### 3.2. Sex-specific effects of chronic DC on serum and liver biomarkers

Several serum and tissue biomarkers were examined to explore the role of DC on glucose, lipid, and hepatic metabolism (Table 1). Interestingly, DC could not change any glycemic parameter (glucose and TyG index) in female and male mice. Likewise, TG and glycerol (an indirect marker of lipolysis) remained similar to the C group in female and male mice receiving DC (DC and F/DC groups). DC did not alter the serum markers of liver injury ALT, AST, and GGT in females. Still, in males, both DC and F reduced ALT compared to the C group (DC −30.1%, F −53.2%, and F/DC −32.1%, p<0.0001), and this parameter was influenced by F and F/DC interaction (p<0.0001, two-way ANOVA). Finally, bilirubin did not change, and albumin levels decreased in male DC, F, and F/DC groups compared to the C group and were influenced by DC and its interaction with F (p<0.0001, two-way ANOVA).

**Table 1.** Serum and hepatic biochemistry.

| Parameters | Female |  |  |  | Male |  |  |  |
| --- | --- | --- | --- | --- | --- | --- | --- | --- |
|  | C | DC | F | F/DC | C | DC | F | F/DC |
| Glucose, mg/dL | 135.0 ± 3.6 | 133.1 ± 5.2 | 148.0 ± 4.5 | 133.4 ± 4.2 | 151.8 ± 14.4 | 146.7 ± 24.6 | 128.1 ± 22.6 | 154.5 ± 53.4 |
| TyG index | 8.35 ± 0.54 | 8.42 ± 0.45 | 8.86 ± 0.26 | 8.58 ± 0.37 | 4.89 ± 0.22 | 4.76 ± 0.22 | 4.61 ± 0.33 | 4.76 ± 0.24 |
| TG, mg/dL | 70.7 ± 32.6 | 75.4 ± 37.4 | 97.6 ± 22.4 | 84.8 ± 31.4 | 119.9 ± 42.5 | 107.5 ± 37.3 | 93.0 ± 40.0 | 105.8 ± 36.0 |
| Glycerol, mg/dL | 51.4 ± 14.0 | 42.4 ± 10.8 | 45.0 ± 5.6 | 53.2 ± 30.4 | 493.4 ± 182.5 | 320.5 ± 286.1 | 371.0 ± 271.4 | 429.1 ± 336.4 |
| ALT, IU | 4.63 ± 1.67 | 4.39 ± 1.14 | 4.31 ± 2.55 | 3.05 ± 2.16 | 7.81 ± 0.96 | 5.46 ± 0.95<sup>a</sup> | 3.66 ± 0.67<sup>a,b</sup> | 5.30 ± 1.80<sup>a</sup> |
| AST, IU | 7.86 ± 1.67 | 7.49 ± 1.63 | 6.57 ± 1.65 | 7.90 ± 1.98 | 8.98 ± 3.66 | 7.97 ± 1.37 | 8.11 ± 1.65 | 7.81 ± 2.76 |
| GGT, IU | 4.01 ± 0.91 | 3.83 ± 0.82 | 4.46 ± 2.08 | 3.44 ± 2.03 | 3.93 ± 0.63 | 3.25 ± 1.68 | 3.50 ± 0.98 | 3.01 ± 1.82 |
| Albumin, mg/dL | 2.31 ± 0.26 | 2.13 ± 0.25 | 2.13 ± 0.27 | 1.94 ± 0.26 | 1.74 ± 0.11 | 1.62 ± 0.07<sup>a</sup> | 1.49 ± 0.07<sup>a,b</sup> | 1.42 ± 0.09<sup>a</sup> |
| Bilirubin, mg/dL | 0.48 ± 0.15 | 0.54 ± 0.16 | 0.51 ± 0.15 | 0.52 ± 0.12 | 0.93 ± 0.59 | 0.86 ± 0.10 | 0.52 ± 0.50 | 0.57 ± 0.47 |

Data are expressed as mean ± S. D. When indicated, p<0.05, [a] C, [b] DC, [c] F, two-way ANOVA, and Holm-Sidak post hoc test. Abbreviations: C, control. DC, sodium deoxycholate. F, fructose. TyG, triglyceride-glucose index. TG, triglyceride. ALT, alanine aminotransferase. AST, aspartate aminotransferase. GGT, gamma-glutamyl transferase (GGT).

### 3.3. DC did not change liver mass, hepatocyte size, or protein expression

Fructose increased liver mass in female F and F/DC groups compared to the C group but not in male mice (Fig. 2A). Then, we assessed the number density of hepatocyte nuclei and the volume-weighted mean nuclear volume since they are indirect and direct measurements, respectively, of hepatocyte hypertrophy (Fig. 2B-C). In female mice, the number density of nuclei was not changed by DC (Fig. 2B), while a greater nuclei volume was seen in the F group (+53.0%, p<0.0001, Fig. 2C-D). In male mice, the number density of hepatocytes and the nuclei volume showed no change, as shown in Fig. 2B-D. The analysis of hepatic TG and glycerol revealed no impact of chronic DC use on tissue metabolism in either female or male mice (Fig. 3A-B). In females, F and F/DC groups showed a higher degree of hepatic fat deposition compared to the group C group (F 305.3%, F/DC 351.4%, p<0.005, Fig. 3C), evidenced by liver photomicrographs in Fig. 3F. On the opposite way, male’s liver showed no change in fat deposition (Figs. 3C, G). Regarding protein expression, FXR, a major regulator of bile acids, did not change its expression by DC in females and males (Fig. 3D), similar to FGF21, which is involved in free fatty acid β-oxidation (Fig. 3E).

**Figure 2.**
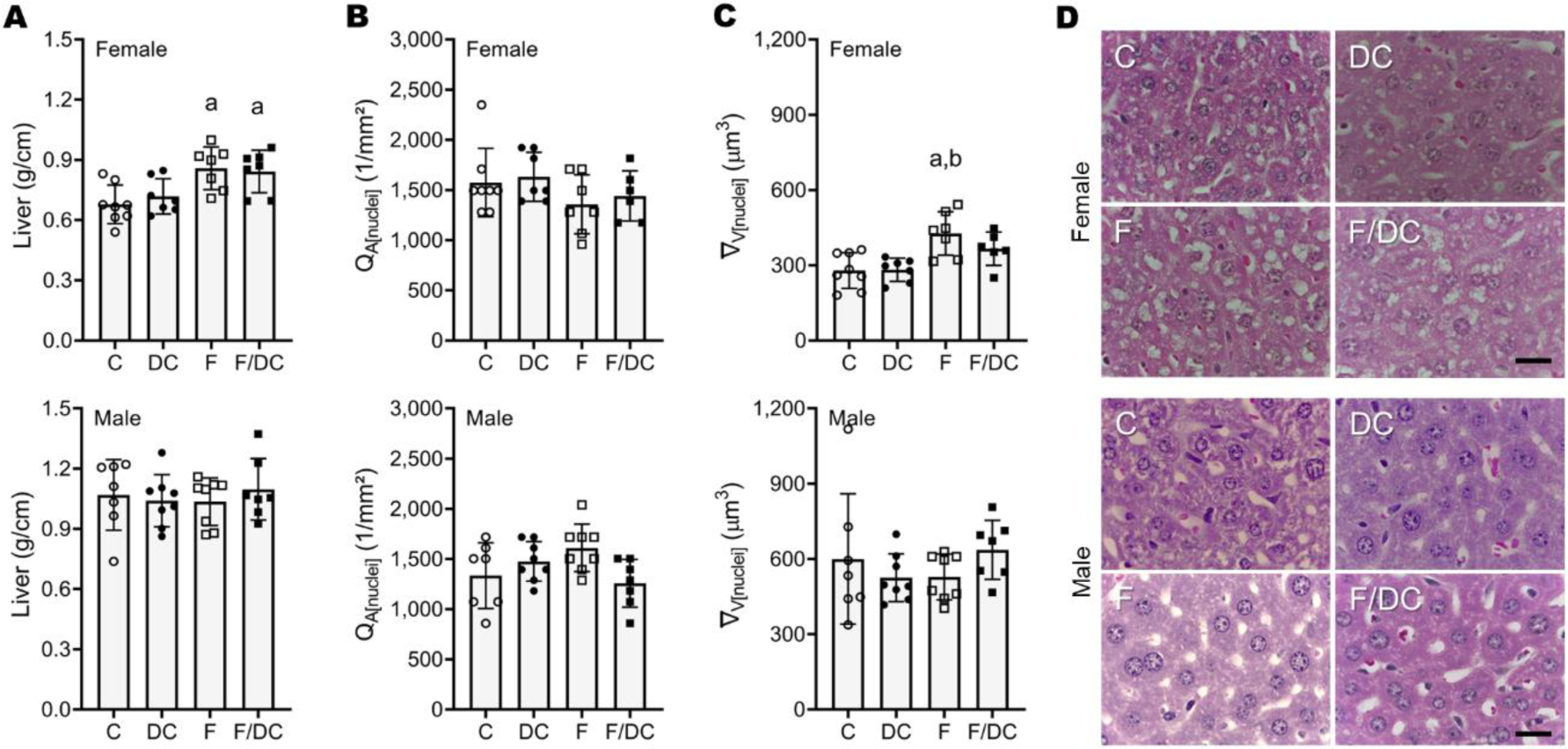
– Liver mass and morphology in female and male Swiss mice. **A**, Liver mass. **B**, Number density of hepatocytes (*Q*_*A*[*nuclei*]_), and **C**, Volume-weighted mean nuclear volume 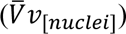 of hepatocytes. **D**, Representative liver photomicrographs (HE stain). Data are shown as mean ± S. D., p<0.05, [a] vs. C, [b] vs. DC. Abbreviations: C, control; DC, sodium deoxycholate; F, fructose; F/DC, fructose + sodium deoxycholate. Bar = 20 µm.

**Figure 3.**
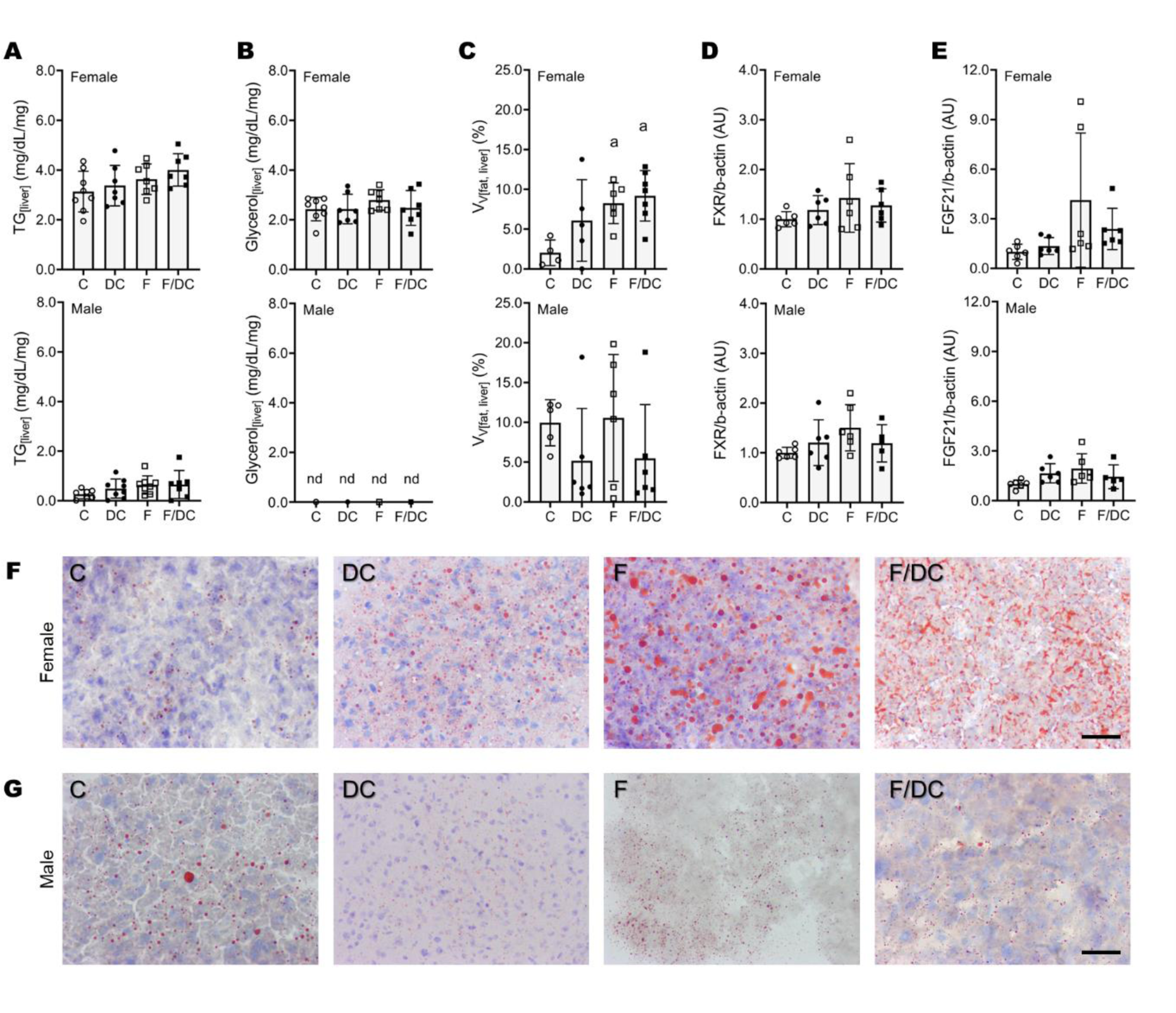
– Liver metabolism and protein expression in female and male Swiss mice. **A**, Hepatic triglycerides **B**, Hepatic glycerol. **C**, Volume density (*Vv*_[*lipid droplets*]_) of hepatic fat. **D**, FXR relative protein expression. **E**, FGF21 relative protein expression. **F**, Fat deposition in female mice (Oil red O stain). **G**, Fat deposition in male mice (Oil red O stain). Data are shown as mean ± S. D., p<0.05, [a] vs. C. Abbreviations: C, control; DC, sodium deoxycholate; F, fructose; F/DC, fructose + sodium deoxycholate; TG, triglycerides; FXR, farnesoid X receptor; FGF21, fibroblast growth factor 21; AU, arbitrary units. Bar = 60 µm.

## 4. Discussion

In Humans, DC promotes the local lysis of adipocyte cell membrane, triggering an inflammatory cascade [28] that takes approximately 28 days to resolve completely [11]. Some side effects are observed at the application site, such as pain, swelling, numbness, bruising, induration, nodules, and itching [29–31]. Also, DC-induced adipocyte destruction reduces body measurements but does not promote BW loss [8]. In the present study, we successfully reproduced DC-induced riWAT inflammation without BW loss in female and male Swiss mice.

Furthermore, DC mesotherapy in humans is unable to permanently alter serum free fatty acids, total cholesterol (TC), low-density lipoprotein (LDL), high-density lipoprotein (HDL), TG, and pro-inflammatory cytokines [9, 32] Our study could also not show long-term metabolic impairments in glucose, lipolysis, and hepatic enzymes. The only exception was ALT and albumin in males, whose levels were decreased in DC and F/DC groups, suggesting a sex-specific effect regardless of fructose consumption. In humans, low ALT levels are associated with a higher risk of mortality [33], and hypoalbuminemia is an essential marker of the severity of body inflammation and physiological stress [33].

Finally, we applied stereological tools to investigate if DC would change liver morphology and quantified the relative protein expression of FXR and FGF21 by western blot, and we failed to demonstrate an adverse role of DC on these parameters. To date, it is the first study investigating the role of chronic DC use in mesotherapy in a mouse model focusing on hepatic outcomes and comparing sexes, which limits our discussion due to the lack of additional evidence from other research groups.

DC-induced adipocyte membrane rupture releases a massive amount of TG into the interstitium, and macrophages are recruited to remove these lipids [13], as evidenced by foam cells (macrophages) in crown-like structures surrounding adipocyte cells [34]. Hypothetically, TG phagocytosis by macrophages might prevent TG from freely entering the bloodstream [35]. However, to what extent macrophages can deal with this TG overload, especially in the long term after successive DC injections, is still unknown. The question remains regarding where these lipids go and if macrophages can phagocyte all of it. Thus, if DC is meant to be used for body contouring, it is essential to establish safety maximum doses that can reduce subcutaneous fat without leading to massive WAT TG release.

In recent years, it has been discovered that bile acids such as DC can bind to multiple tissue receptors and act as metabolic regulators of pathways related to glucose, lipids, and amino acids metabolism, homeostasis maintenance, and gut microbiota, which explain their implication in several metabolic disorders [17, 20]. In the liver, DC binds to FXR, which regulates glycolipid metabolism and endothelial function, which are enrolled in cardiovascular disease risk [17]. In turn, FXR activation reduces circulating DC [36] because it acts as a bile acid sensor in the enterohepatic circulation, and it maintains bile acid homeostasis by modulating the expression of genes enrolled in bile acid synthesis, transport, and excretion [21].

Glycerol is an indirect biomarker of lipolysis that can be transported from the adipose tissue into the liver to be stored as TG. In addition, the liver can store TG under stress conditions such as abnormal lipid metabolism, high-fat high-carbohydrate intake, and obesity [37]. We did not observe changes in glycerol and TG levels induced by DC, which corroborates with FGF21 protein expression. Among its functions, FGF21 stimulates the β-oxidation of free fatty acids and suppresses the formation of TG and adipose tissue lipolysis, reducing TG’s intrahepatic and serum content [22]. Although we have shown that fructose feeding led to mild fat deposition in the liver of female mice, corroborating an unbalanced diet intake, it was not sufficient to modulate hepatic FGF21 protein expression.

Non-surgical fat reduction ranks fourth among men’s most common non-surgical procedures and fifth in women [38]. Firstly, it surprisingly shows that the search for aesthetic procedures for fat reduction is not limited to women, and secondly, it highlights the relevance of performing studies investigating both sexes. As in humans, studies in animal models have already shown sex-specific effects, mediated by sex hormones, on adipocyte development, adipogenesis, lipogenesis, lipolysis, and insulin sensitivity [39]. Thus, sex hormones may impact the body’s response to DC mesotherapy, as well as to other substances that may be under investigation for body fat reduction.

We chose the Swiss mice for DC mesotherapy since the healthy mice have a large subcutaneous fat compartment at the inguinal region [40]. It is easily accessible for the administration of substances, and we have succeeded in DC administration at the riWAT. We also offered fructose to investigate the impact of DC mesotherapy on an organism fed with an unhealthy diet since fructose administration has been widely used to induce obesity and the metabolic syndrome phenotype in animal models [41]. In Swiss mice, most studies offer fructose in drinking water, ranging from 15% to 30% [42–44]. According to Johnson et al., 10% fructose would be sufficient to induce hypertension and microvascular renal changes [45]. Based on this evidence, we offered an intermediate dose of 20% fructose in drinking water for 12 weeks. Unfortunately, we did not find expressive BW gain and metabolic changes in fructose-feed mice of both sexes, which may be a limitation of this study.

In conclusion, DC did not impact BW or adipose tissue mass. However, subcutaneous DC injections promoted adipocyte lysis and inflammation, evidenced by riWAT hemorrhage, foam cells, and tissue fibrosis. Surprisingly, the chronic usage of DC failed to impair glucose and hepatic metabolism, hepatic morphology, and protein expression in both sexes, even in mice under fructose feeding. Thus, in the present mice model, chronic DC mesotherapy proved safe for liver metabolism. Overall, care must be taken in extrapolating this data to humans since further studies are required to prove its safety in other body systems such as the heart and kidneys.

## Acknowledgments

The authors are thankful to Priscila Rodriguez Câmara and Clara Ana Santos Monteiro for their technical assistance. Authors are also thankful for the student scholarships provided by the Coordination for the Improvement of Higher Education Personnel (Coordenacao de Aperfeicoamento de Pessoal de Nivel Superior/CAPES) [Finance Code 001] and the Pro-Reitoria de Pesquisa, Pós-Graduação e Inovação (PROPPI) of Universidade Federal Fluminense.

